# Japanese encephalitis virus capsid protein interacts with non-lipidated MAP1LC3 on replication membranes and lipid droplets

**DOI:** 10.1101/2020.08.04.237248

**Authors:** Riya Sarkar, Kiran Bala Sharma, Anita Kumari, Shailendra Asthana, Manjula Kalia

## Abstract

Studies have shown that Japanese encephalitis virus (JEV), replicates on ER derived membranes that are marked by autophagosome negative non-lipidated MAP1LC3 (LC3-I). Depletion of LC3 exerts a profound inhibition on virus replication and egress. Here, we further characterize the role of LC3 in JEV replication, and through immunofluorescence and immunoprecipitation show that LC3-I interacts with the virus capsid protein in infected cells. This association was observed on capsid localized to both the replication complex and lipid droplets (LDs). JEV infection decreased the number of LDs per cell indicating a link between lipid metabolism and virus replication. This capsid-LC3 interaction was independent of the autophagy adaptor protein p62/SQSTM1. Further, no association of capsid was seen with the GABARAP protein family, suggesting that this interaction was specific for LC3. High resolution protein-protein docking studies identified a putative LC3-interacting region (LIR) in capsid, _56_FTAL_59_, and other key residues that could mediate a direct interaction between the two proteins.

## Introduction

Flavivirus genome replication takes place in virus induced specialised intracellular membranous structures described as convoluted membranes (CMs) and vesicle packets (VPs). These originate from the ER and are the sites for polyprotein processing/translation and viral RNA replication respectively [1–3]. The VPs are composed of both viral and host proteins, and confine viral RNA replication to specific cytoplasmic locations. This serves the dual purpose of shielding the viral RNA from the host innate immune system and concentrating the components required for replication [4–7].

Replication complex biogenesis begins with the modification of the lipid composition and protein interactions on the ER membranes. The flavivirus replication complex is composed of viral proteins NS4a, NS1, NS2a, NS5 and viral dsRNA replicative intermediate [6, 8, 9]. The complex is connected to the cytosol via a pore, which acts as a passage for transport of nucleotides and other factors required for RNA replication. Structural proteins precursor to-membrane (prM), capsid (C) and envelope (E) are not a part of these replication complexes, but the formation of nucleocapsid takes place near the RNA exit site [2, 10]. Viruses hijack a diverse array of host proteins to induce the formation of these replication complexes [11–14].

Flaviviruses such as Dengue virus (DENV) give rise to negative invaginated vesicles towards the ER lumen [2], while Hepatitis C virus (HCV) forms double membrane vesicles of positive curvature towards the cytoplasm. HCV shows a unique complex membrane rearrangement known as membranous web in close association with lipid droplets (LDs) which helps in viral replication and virus assembly [15].

LDs are cytoplasmic organelles enveloped within a single phospholipid membrane and store neutral lipids (mainly triacylglycerol and cholestrol esters). Their numbers and size in different cell types is highly variable and dynamic. LDs play a crucial role in lipid metabolism and impact cellular homeostasis and processes such as signaling, immune responses and pathogen infection [16]. LDs have been described as platforms for HCV assembly, with the capsid localizing to the lipid droplets and subsequent recruitment of NS5A which engages virus replication complexes to LD-associated membranes [17, 18]. All the other flavivirus capsid proteins-DENV, Zika virus (ZIKV), West nile virus (WNV) and JEV have been reported to localize on LDs and this association is crucial for virus particle formation [19–22]. Virions are assembled in close proximity to the ER and LDs and bud into the ER-lumen for envelopment followed by transport through the secretory pathway [23]. LDs can also provide energy for virus replication [16, 24].

Microtubule-associated protein 1 light chain 3 (MAP1LC3, and henceforth LC3) is an ubiquitin-like protein and its lipidated form (LC3-II) is a defining characteristic of autophagosomes [25]. However, the non-lipidated LC3 (LC3-I) also has autophagy independent roles and associates with ER associated degradation (ERAD) protein enriched membranes [26–28]. The presence of non-lipidated LC3 as a part of the virus replication complex was first reported for mouse hepatitis coronavirus (MHV) [29], and subsequently for the equine arteritis virus (EAV) and JEV [27, 30]. These studies also showed that depletion of LC3 resulted in a significant decrease in virus replication, validating its role as an essential host factor. Other viruses such as Coxsackievirus, Polio virus (PV), DENV and ZIKV can also utilize autophagy independent LC3 as a membrane source for replication [28, 31]. A recent study has shown the association of Influenza virus (IAV) capsid protein with LC3 [32].

Here we demonstrate that the JEV capsid protein associates with LC3-I in infected cells. This association was also observed on lipid-enriched membranes and is likely to be essential for ribonucleoprotein and subsequent infectious virus particle formation. The number of lipid droplets decreased significantly in JEV infected cells highlighting a link of virus replication with lipid droplet metabolism. The capsid-LC3 interaction was independent of the key autophagy adaptor protein SQSTM1. A detailed molecular modelling study identified a putative LC3-interacting region in the capsid protein, and key residues that are likely to be involved in LC3-capsid interaction.

## Results

### Subcellular localization of JEV capsid in infected cells

Our earlier studies have demonstrated the immunofluorescence staining pattern of JEV NS proteins-NS1, NS3 and NS5 in infected Neuro2a, Huh7 and MEFs [12, 27]. These were majorly observed in an irregular-shaped punctate network that extended throughout the cell and likely represents CMs. The dsRNA staining is seen as discrete puncta with significant overlap with the NS proteins [11, 12, 27]. Here we have analysed capsid staining in JEV infected cells (Fig 1). In accordance with published literature we observed capsid in close juxtaposition with replication complexes, marked here by the NS1 protein in infected HeLa and Huh7 cells (Fig 1A, C). In several images extensive overlap was also observed between NS1 and capsid which could be because of the limitation of confocal (light) microscopy to resolve these structures (Fig 1C). However, in high-resolution SIM images, NS1 and capsid were clearly seen in close proximity but distinct (Fig 1D). Nuclear localization of capsid was also observed in both cell type (Fig 1A, B, E extreme right panels). To check for LD localization, JEV infected cells were incubated with BODIPY which stains neutral lipid. While only a few LDs were seen in HeLa cells, Huh7 hepatocytes showed an abundant distribution. Capsid was observed on LDs in both cell types (Fig 1B, E). The capsid decorated LDs localized both to the cytosol (Fig 1B, E, centre panels), and to the nucleus (Fig 1B, E, right panels) of infected cells.

**Figure 1:**
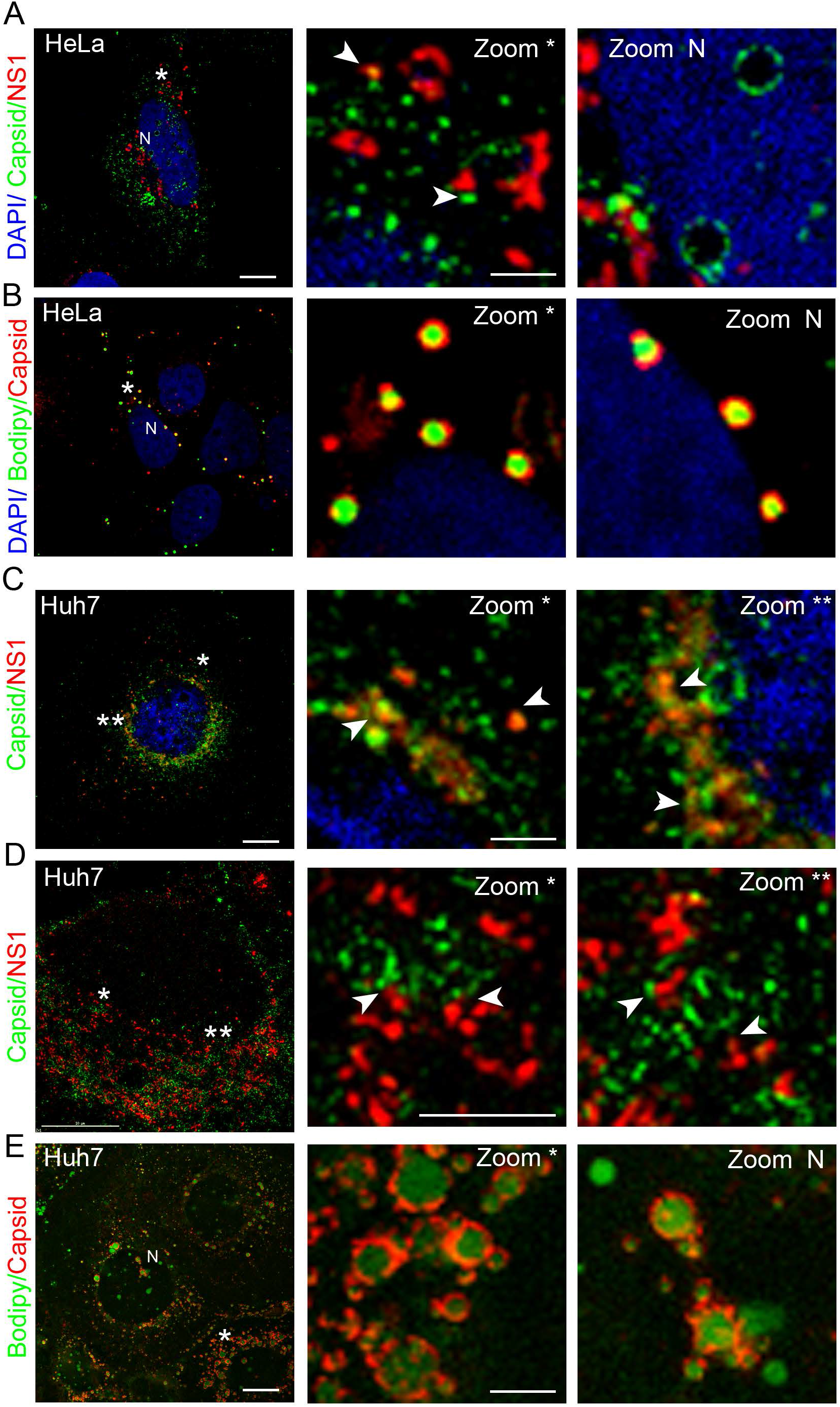
JEV capsid localization in infected HeLa and Huh7 cells. HeLa (A-B), and Huh7 cells (C-E), were infected with JEV (5MOI, 24 h). Cells were stained with capsid (green) and NS1 (red) antibodies (A, C, D); or with Bodipy (green) and capsid (red) (B, E). Panel D is a SIM image, while the others are confocal images. Middle and right panels show a magnified view of the region marked by */ **/ N (nucleus). Arrows indicate areas of colocalization and close proxomity between NS1 & capsid staining. Scale bar, 10 μm (left panel) and 2 μm (middle & right panels). Images are representative of three or more independent experiments.

We next checked if virus infection changed the distribution and number of LDs in the cell. Mock and JEV-infected Huh7 cells were incubated with BODIPY (Fig 2A-C), and as expected capsid was observed to be strongly associated with LDs in infected cells (Fig 2B), while NS1 was in close proximity but not associated with these structures (Fig 2C). Interestingly, while no change was discernible in the distribution or size of LDs, their number decreased significantly in JEV infected cells (Fig 2D). This suggests that virus infection impacts the cellular lipid metabolism.

**Figure 2:**
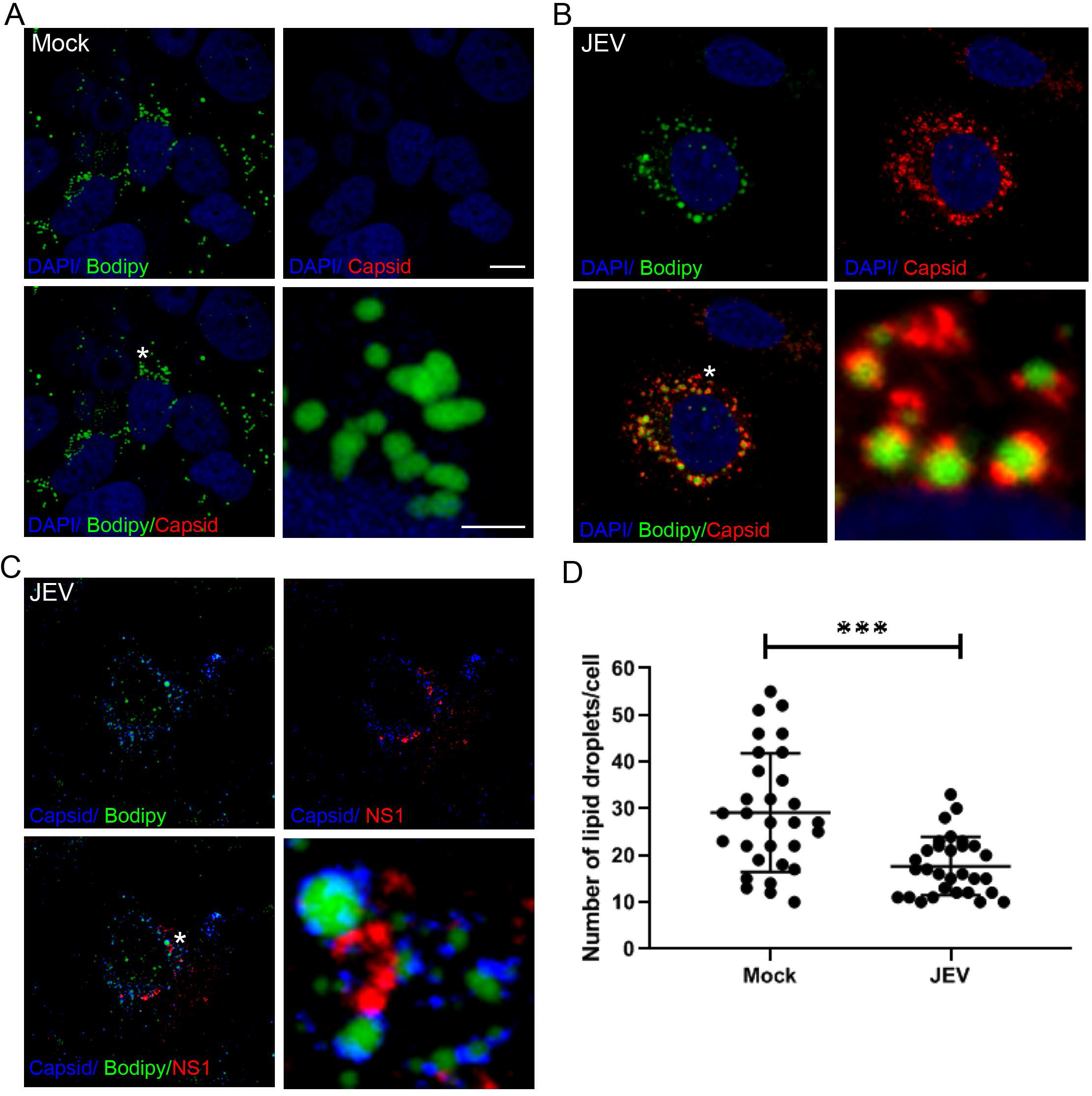
JEV capsid accumulates around lipid droplets in infected Huh7 cells. (A-C) Huh7 cells were either mock infected (A), or infected with JEV (5MOI, 24 h), and stained using Bodipy (green) and capsid (A, B) or NS1 (C) antibodies (red). Nuclei were visualized by DAPI staining (blue). Colour merged images are shown in the lower left panels. Lower right panels show a magnified view of the region corresponding to the asterisk (*). Scale bar, 10 μm and 2 μm (lower right panels). (D) The number of lipid droplets in mock and JEV infected cells was quantified from 30 cells across three coverslips using Image J. Values are shown as mean ± SD. Student’s t-test was used for comparing data from mock and JEV infected cells. ***, p<0.001.

### JEV capsid interacts with LC3-I in WT and *atg5−/−* MEFs

The LC3 protein is translated as a full-length precursor designated as proLC3 which undergoes cleavage at a highly conserved Gly120 residue resulting in the formation of LC3-I, which can then undergo lipidation to give rise to the autophagosome incorporated LC3-II [33]. On western blots LC3-I (~16-18 kDa) can be seen as an upper band, while LC3-II (~14-16 kDa) displays faster electrophoretic mobility and is visible as a lower band (Fig 3C, input sample). LC3-I is predominantly cytosolic, however it has been shown to associate with virus replication complexes for MHV, EAV and JEV. These complexes are visible through immunofluorescence staining using antibodies against LC3 in both autophagy competent and deficient cells [27, 29, 30]. We have shown that in Neuro2a, WT, and autophagy deficient (*atg5−/−*) MEFs, virus replication complexes marked by NS1 show colocalization with autophagosome independent LC3-I and EDEM1 [27]. Depletion of LC3 by siRNA results in reduced virus replication validating its role as an essential host-factor [27, 29, 30]. However, immunoprecipitation of JEV-NS1 from infected cells did not pull down any LC3 (our unpublished observations). In this study we checked the localization of the JEV capsid protein with endogenous LC3 in infected WT and *atg5−/−* MEFs. Capsid protein staining was seen as distinct large foci and showed strong co-localization with LC3 in both cell types (Fig 3A, B). The Pearson’s correlation coefficent between capsid and LC3 in WT and *atg5−/−* MEFs was 0.65 and 0.79 respectively. Our earlier studies have established that in autophagy competent cells (Neuro2a & WT MEFs) these replication complex & LC3 positive structures are not autophagosomes, as they are negative for LAMP-1, Lysotracker Red and do not overlap with GFP-LC3 [27]. On the other hand in autophagy deficient MEF’s only the LC3-I form is observed due to lack of the critical autophagy protein ATG5 that is essential for lipidation of LC3-I to LC3-II [34] (Fig 3D, input). Significant overlap of LC3 with LD localized capsid was also observed in *atg5−/−* MEFs (Fig 4).

**Figure 3:**
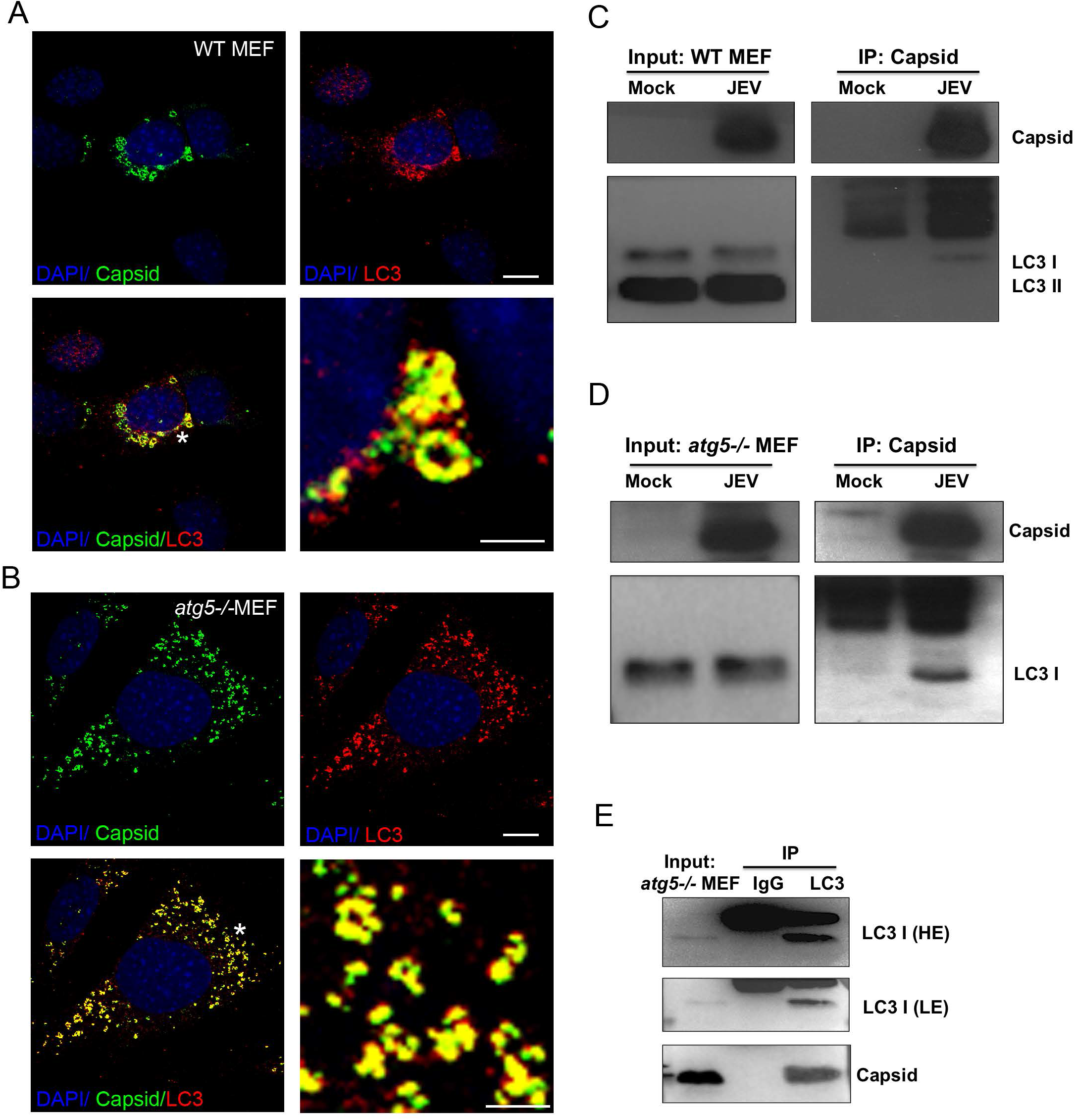
JEV capsid protein colocalizes with endogenous LC3 in infected WT and *atg5−/−* MEFs. (A, B) JEV infected (5 MOI, 24 hpi) WT (A) and *atg5−/−* MEFs (B) were stained with capsid (green) and LC3 (red) antibodies. Nuclei were visualized by DAPI staining (blue). Colour merged images are shown in the lower left panels. Lower right panels show a magnified view of the region corresponding to the asterisk (*). Scale bar, 10 μm and 2 μm (lower right panels). Images are representative of three or more independent experiments. (C-D) Mock/JEV infected (5 MOI, 24 hpi) WT (C) and *atg5−/−* MEFs (D) were lysed and immunoprecipitation was performed using capsid antibody. Western blots showing capsid and LC3 proteins in input (left panel) and IP samples. (E) JEV infected *atg5−/−* MEF lysates were immunoprecipitated using rabbit IgG or LC3 antibodies, and blotted for LC3 and capsid. The blots are representative of two independent experiments.

**Figure 4:**
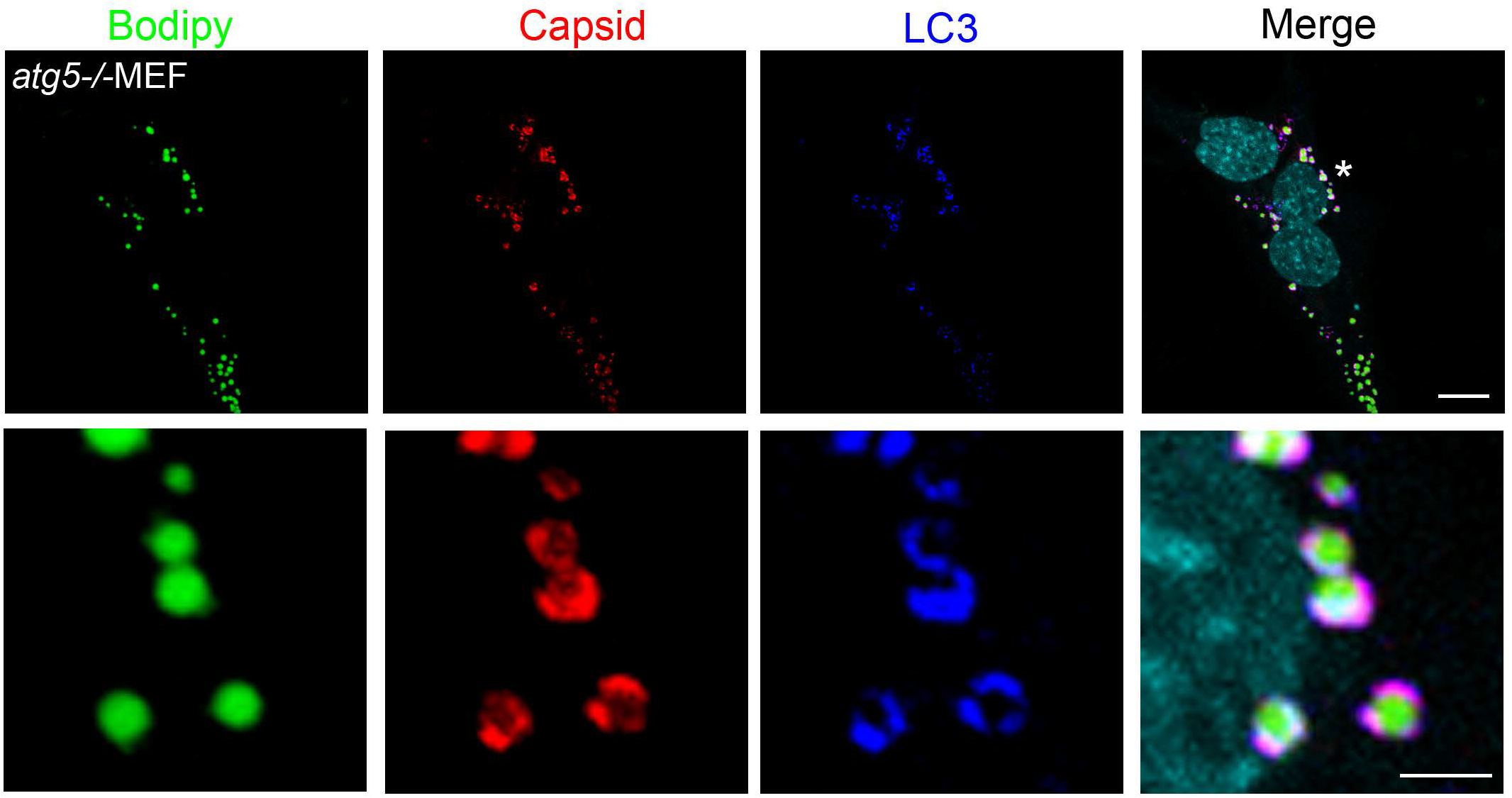
JEV capsid protein localizes with lipid droplets and LC3-I in autophagy deficient MEFs. JEV infected (5 MOI, 24 hpi) *atg5−/−* MEFs were stained with Bodipy (green), capsid (red) and LC3 (blue) antibodies. Nuclei were visualized by DAPI staining (cyan in merge image). Lower panels show a magnified view of the region corresponding to the asterisk (*). Scale bar, 10 μm (upper panel) and 2 μm (lower panel).

We further validated this interaction by immunoprecipiatation studies. Pull down of capsid protein from JEV infected WT MEF’s, also brought down LC3-I and not LC3-II (Fig 3C). We also checked this in autophagy deficient cells and observed that immunoprecipiation with capsid brought down LC3-I protein in these cells (Fig 3D). We further performed an inverse IP from JEV infected cells using Rabbit IgG (control) and LC3 antibodies, and were able to pull-down capsid in association with LC3-I from *atg5−/−* MEFs (Fig 3E). These data clearly indicate that LC3-I associates with the JEV capsid protein in infected cells.

### JEV capsid interacts with ectopically-expressed LC3 mutants

To further confirm the specificity of capsid-LC3 interaction, HEK 293 T cells were transfected with Myc-LC3δC22 and Myc-LC3δC22, G120A constructs [33]. The Myc-LC3δC22 has a deletion downstream of Gly 120, and is essentially LC3-I that is capable of lipidation and forming LC3-II. In transfected cells this over-expressed protein can be detected with both Myc and LC3 antibodies through immunofluorescence (Fig 5A, upper left panel), and shows both LC3-I and LC3-II forms on western blots (Fig 5B, input panels). This construct also showed extensive overlap with the capsid protein in infected cells (Fig 5A). Immunoprecipiation experiments using Myc as bait could pull down JEV-capsid from infected cells indicating interaction between the two proteins (Fig 5B).

**Figure 5:**
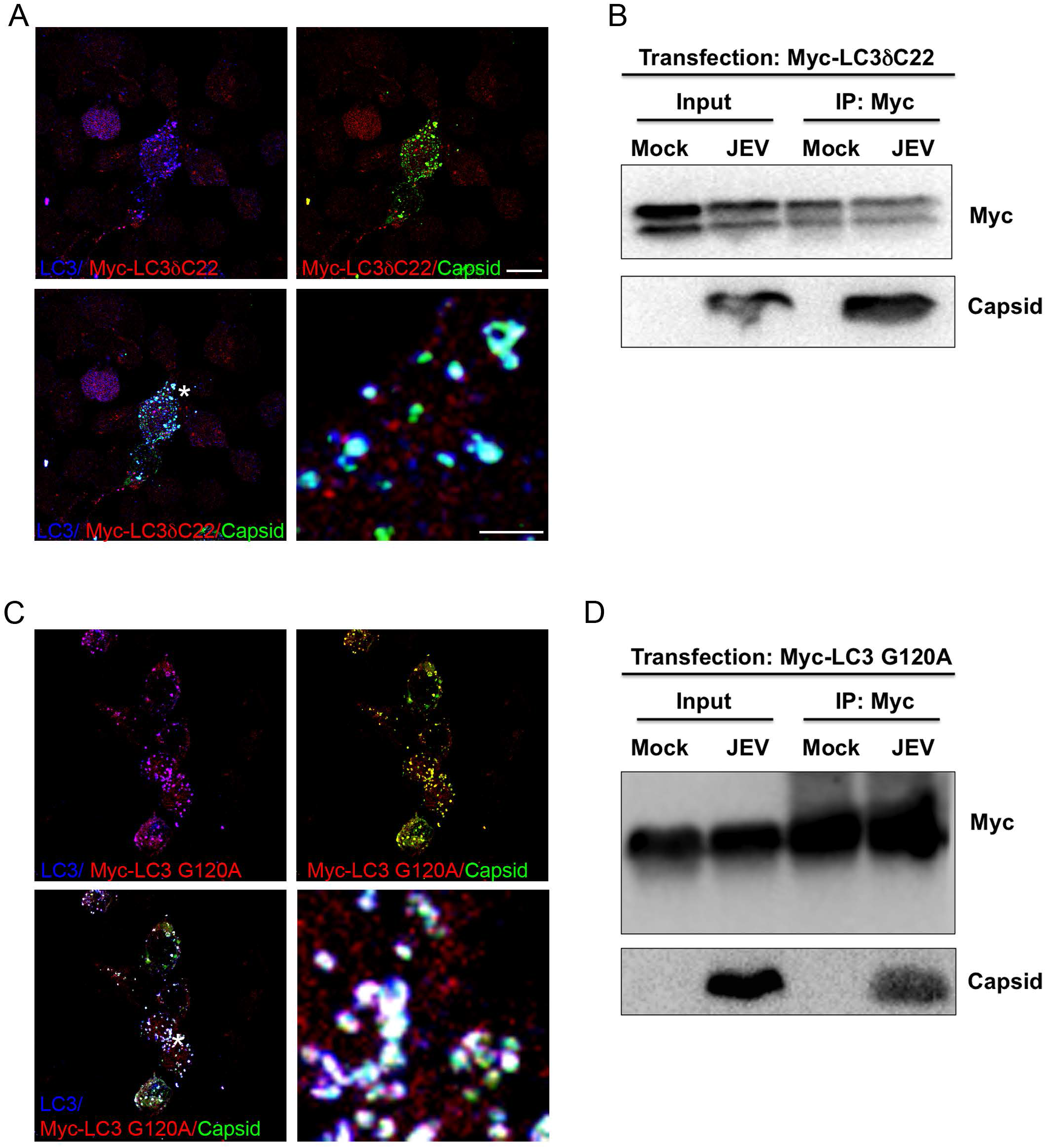
JEV capsid protein colocalizes with ectopically expressed Myc-LC3δC22 and Myc-LC3 G120A in infected HEK 293 T cells. HEK 293T were transfected with Myc-LC3δC22 (A, B) or Myc-LC3 G120A (C, D) and after 24 h were infected with 20MOI JEV for another 24 h. (A, C) Cells were stained with LC3 (blue), Myc (red) and capsid (green) antibodies. Colour merged images are shown in the lower left panels. Lower right panels show a magnified view of the region corresponding to the asterisk (*). Scale bar, 10 μm and 2 μm (lower right panels). (B, D) Transfected and mock/JEV infected cell lysates were immunoprecipiated using Myc antibody and blotted for Myc and capsid. The images and blots are representative of two independent experiments.

In a sound set of experiments, cells were transfected with Myc-LC3δC22, G120A. This construct because of G120A mutation cannot be lipidated and hence exists only in the LC3-I form (Fig 5D, input). The Myc-LC3δC22, G120A also showed extensive co-localization with capsid by immunofluorescence (Fig 5C), and could also immunoprecipiate capsid (Fig 5D). Collectively, these data are indicative of an interaction between JEV-capsid and LC3-I in infected cells.

### LC3 is seen on lipid droplets in association with JEV capsid in infected cells

We next tested if LC3 was present on the lipid droplet localized capsid. Immunofluorescence staining showed that while endogenous LC3 did not localize to lipid droplets in mock-infected cells (Fig 6A), a strong co-localization of caspid-LC3 on lipid droplets was observed in infected cells (Fig 6B). The Pearson’s coefficient of colocalization for capsid-LC3 in infected Huh7 cells was observed to be 0.877.

**Figure 6:**
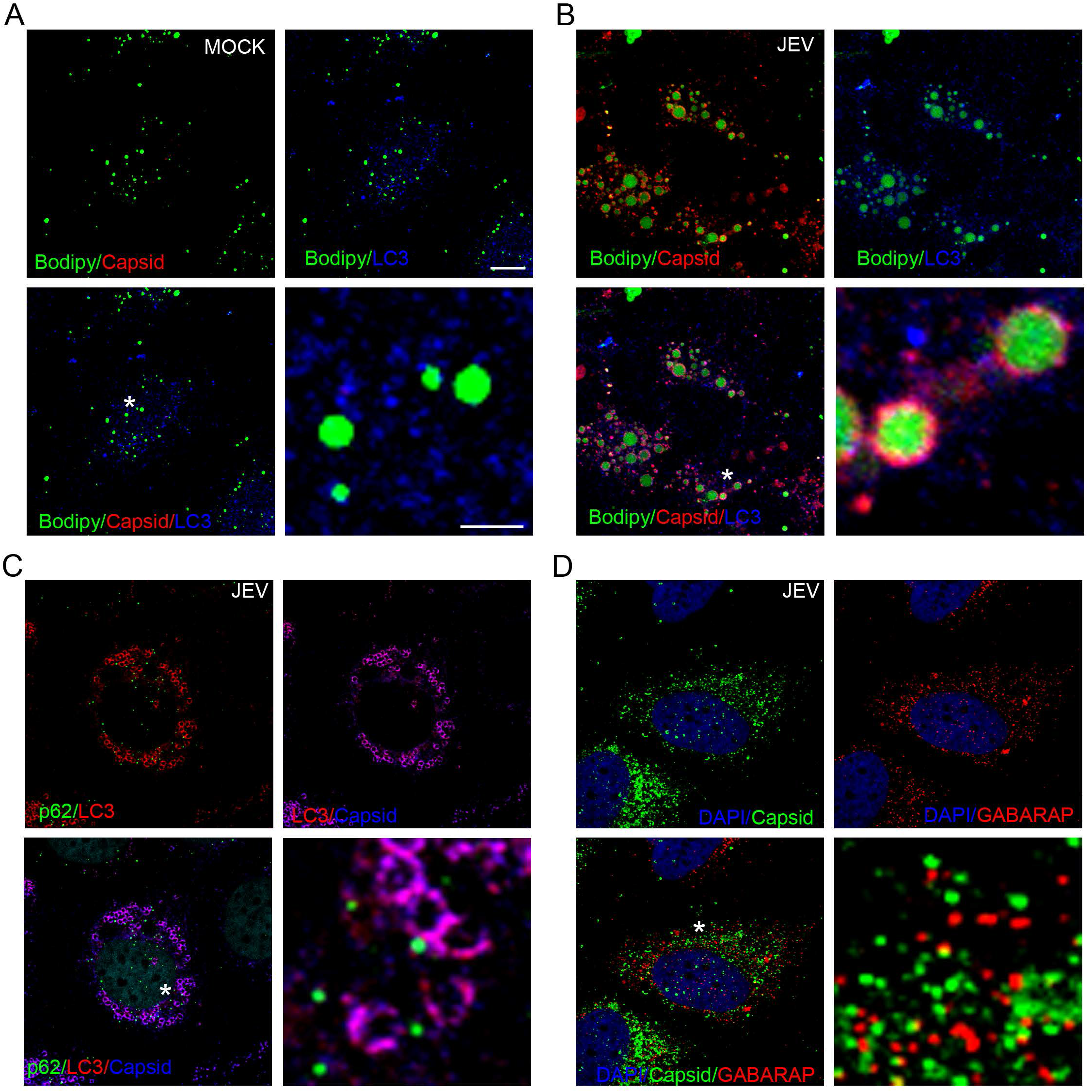
JEV capsid colocalizes with endogenous LC3 around lipid droplets in infected Huh7 cells. Huh7 cells were either mock infected (A), or JEV infected (5MOI) (B-D) for 24 h and were stained using Bodipy (green), capsid (red), LC3 (blue) antibodies (A-B); or with p62 (green), LC3 (red) capsid (blue) antibodies (C); or with capsid (green) and GABARAP (red) antibodies (D). Lower right image in each panel shows a magnified view of the region corresponding to the asterisk (*). Scale bar, 10 μm and 2 μm (lower right image). Images are representative of two or more independent experiments.

Several proteins associate with LC3 through an LC3 interacting region (LIR) motif independent of its lipidation status [35, 36]. SQSTM1/p62 is one such autophagy receptor protein that binds to LC3 via its LIR and targets ubiqutinated cargo to the phagophore [37]. This led us to speculate that p62 could potentially be involved in recruiting capsid to LC3 membranes. However through immunofluorescence staining no overlap between p62 and capsid was detectable (Pearson’s coefficient 0.033), suggesting that the capsid LC3-I interaction takes place independently of p62 (Fig 6C).

γ-aminobutyric acid receptor–associated proteins (GABARAPs) are a second sub-family of Atg8-like proteins in mammalian cells that are involved in bulk sequestration of cytosolic cargo [38]. To check if there is any binding of GABARAPs with capsid, we stained for all three isoforms of GABARAP in JEV infected cells. No overlap was observed between capsid and GABARAP (Pearson’s coefficient 0.032) indicating that the capsid interaction is specific for LC3 (Fig 6D). Collectively our data suggests that the capsid-LC3 interaction is likely to be direct and specific.

### Computational modelling & docking studies for capsid-LC3

To identify the key residues and *hot-spots* involved in capsid-LC3 interactions, the most likely interface site was identified using protein-protein docking approaches [39]. Knowledge of the complete and stable protein structure is a prerequisite for modelling & docking studies. LC3 has been extensively studied in the context of protein-protein interaction [36, 40, 41]. Proteins interact with LC3 through an LC3 interacting motif (LIR motif) and/or LIR like motif [35, 36]. LC3 is highly conserved across species and adopts a well-characterized bilabial fold [42]. Although, the crystal structure of LC3 is reported, a detailed molecular modelling was carried out for loop movements. A similar protocol was executed for the capsid protein also. The most likely poses of the capsid-LC3 complex based on lowest docking energy and number of conformations were generated. These were also compared with those of other LC3 interacting proteins such as p62, FYCO1, FUNDC1 (PDB IDs: 2ZJD, 5D94, 5GMV, respectively).

Both the protein structures displayed an initial structural rearrangement. The overall structural fluctuation of LC3 and capsid was minimal. This was as expected, since the protein structure (template) of LC3 has high resolution and covers more space, while capsid has multiple domains that are poorly connected. Studies have shown that LIR motifs are mainly involved in the interaction with LC3 [36, 40–42]. We analysed the LIR sequences of known LC3 interacting proteins such as PCM1, ATG13, ULK1, FIP200, p62, FYCO1, Influenza M2, Optineurin, ScAtg3, ScAtg19, ScAtg32, NBR1 and BNIP3L/Nix [43], and compared their LIR motif with the structural information of the capsid protein. Our modelling study identified a putative LIR domain in the JEV-capsid protein **_56_FTAL_59_**, which matches well with reported LIR motif (W/F/YxxL/I/V) (Fig 7A-B) [35, 36]. Residues Lys55, Phe56, and Leu59 of capsid interact with LC3 significantly in our model, indicating a possible interaction between capsid and LC3 via this putative LIR motif. This motif was present on the 2^nd^ alpha-helix of capsid, and the Phe56 and Leu59 are highly conserved residues across all *flaviviruses* [44]. The 4^th^ alpha-helix at the C-terminal of capsid (from residues 74 to 98) was also mapped to the LC3 interacting zone (Fig 7C-D).

**Figure 7:**
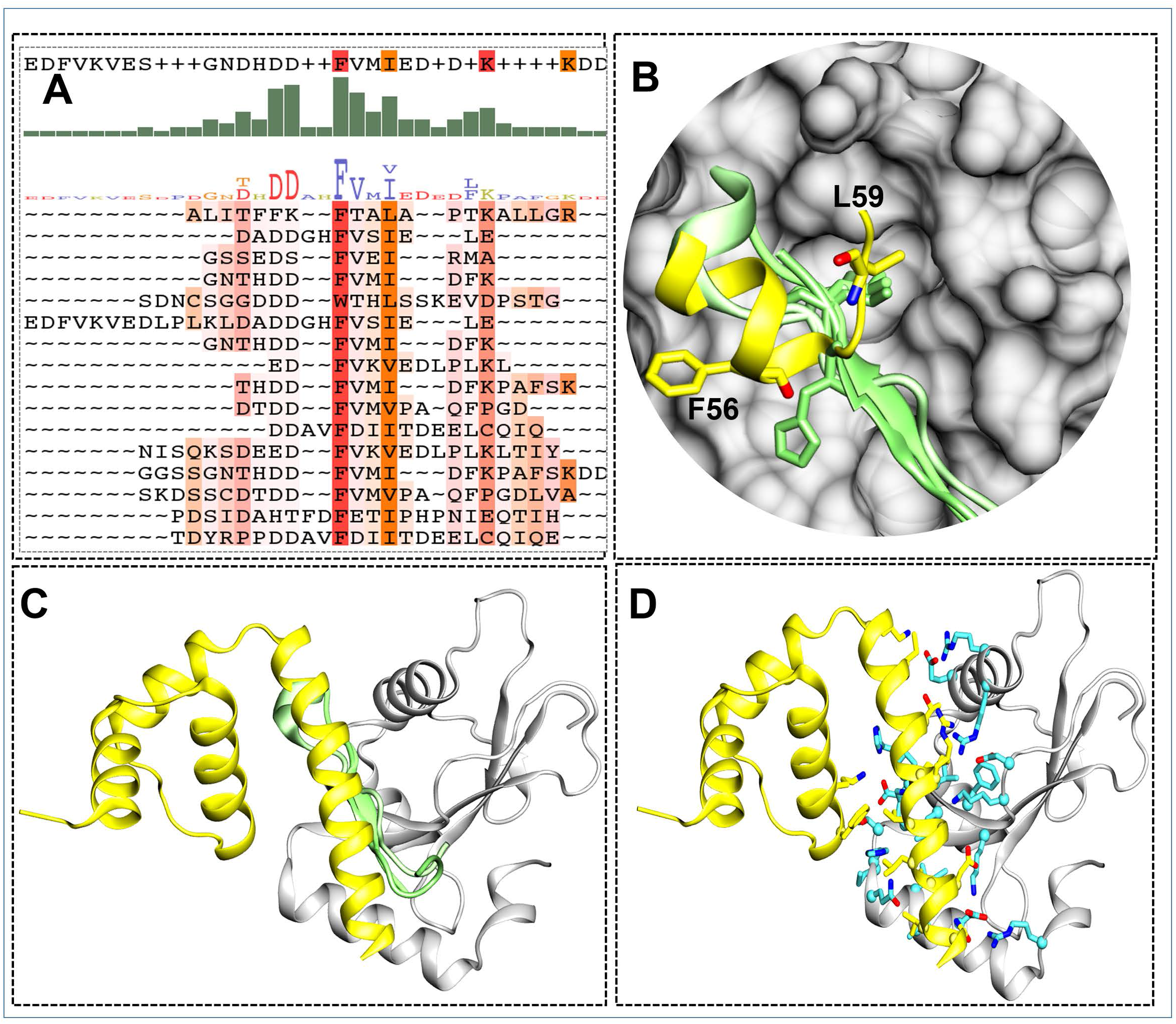
Sequence and structure level characterization of capsid-LC3 interaction. (A) Amino acid sequence alignment and motif generation was performed using sequences of LC3 interacting proteins such as p62, PCM1, ATG13, ULK1, FIP200, FYCO1, Influenza M2, Optineurin, ScAtg3, ScAtg19, ScAtg32, NBR1, BNIP3L/Nix. The motif and confidence score are shown. (B, C and D) The structural alignment of capsid (in yellow) and other LC3 interacting protein structures (PDB-IDs: 2ZJD, 5D94, 5GMV) in green are used. The aligned residue Leu59 is highlighted in B, which matched nicely with the Leu in the LIR motif of known LC3 interactors.

From the lowest docking energy pose of the complex and through guided data from literature, the most likely binding mode of the complex was selected for further quantitative analysis. Three types of interactions were observed at the interface of capsid-LC3, hydrogen bonding (HB), hydrophobic (HpH) contacts, and Pi–Pi interactions. The most frequent HBs formed between the LC3@Glu62:Capsid@Lys74=2.80Å, Leu53:Lys85=2.73Å, Arg11:Asn96=2.61Å, Asp19:Asn96=2.93Å, His57:Lys55=2.92Å, Asp56:Lys55=2.81Å and Thr29:Lys55=2.76Å, Lys49:Lys85=2.73Å. Additional stability was gained through interactions between the HpH residues Leu22, Ile23, Leu53 and Val54 from LC3 and residues Met78, Leu88, Leu91, Ile92, and Val95 from capsid. Of these the capsid Met78 is crucial for virus particle production, and Leu88, Leu91 and Val95 show conservation between groups of strongly similar properties across *flaviviruses* [44]. The basic residues from LC3 are Arg7, His23, Lys26, Lys45, Lys47, His53, Arg65, and Arg66; and from capsid are Lys74, Lys85 and Arg86. The acidic residues of LC3 are Asp19, Asp42, Asp48, Asp56, Glu62, and from capsid only Asp93. Additionally, the Pi–Pi interaction between LC3@His27:Capsid@Phe56=3.8Å also contributed to the higher stability of the complex.

Furthermore, in an effort to dissect these interactions from the docking simulations, total interaction energy between LC3 and capsid was calculated within the framework of Amber-force field description (Fig 8A-B). The residue-wise contribution revealed that Arg11, Asp19, Thr29, Lys49, Leu53 Asp56, His57 and Glu62 of LC3 contribute significantly (<−1.5 kcal/mol: benchmark analysis parameter) either in form of Vander Waal’s forces and/or electrostatically to establish the interactions, with the residues Asp15, Lys45 and Asp52 contributing maximum towards capsid interactions. In case of capsid the significantly contributing residues were Lys55, Lys74, Lys85 and Asn96, with a major role of the three lysines. Studies have shown that the capsid residues Lys85 and Arg86 are crucial for viral RNA packaging [19]. The residue map also shows that the interactions between LC3@Asp19:capsid@Asn96 and LC3@Asp56:capsid@Lys55 may also be contributing significantly.

**Figure 8:**
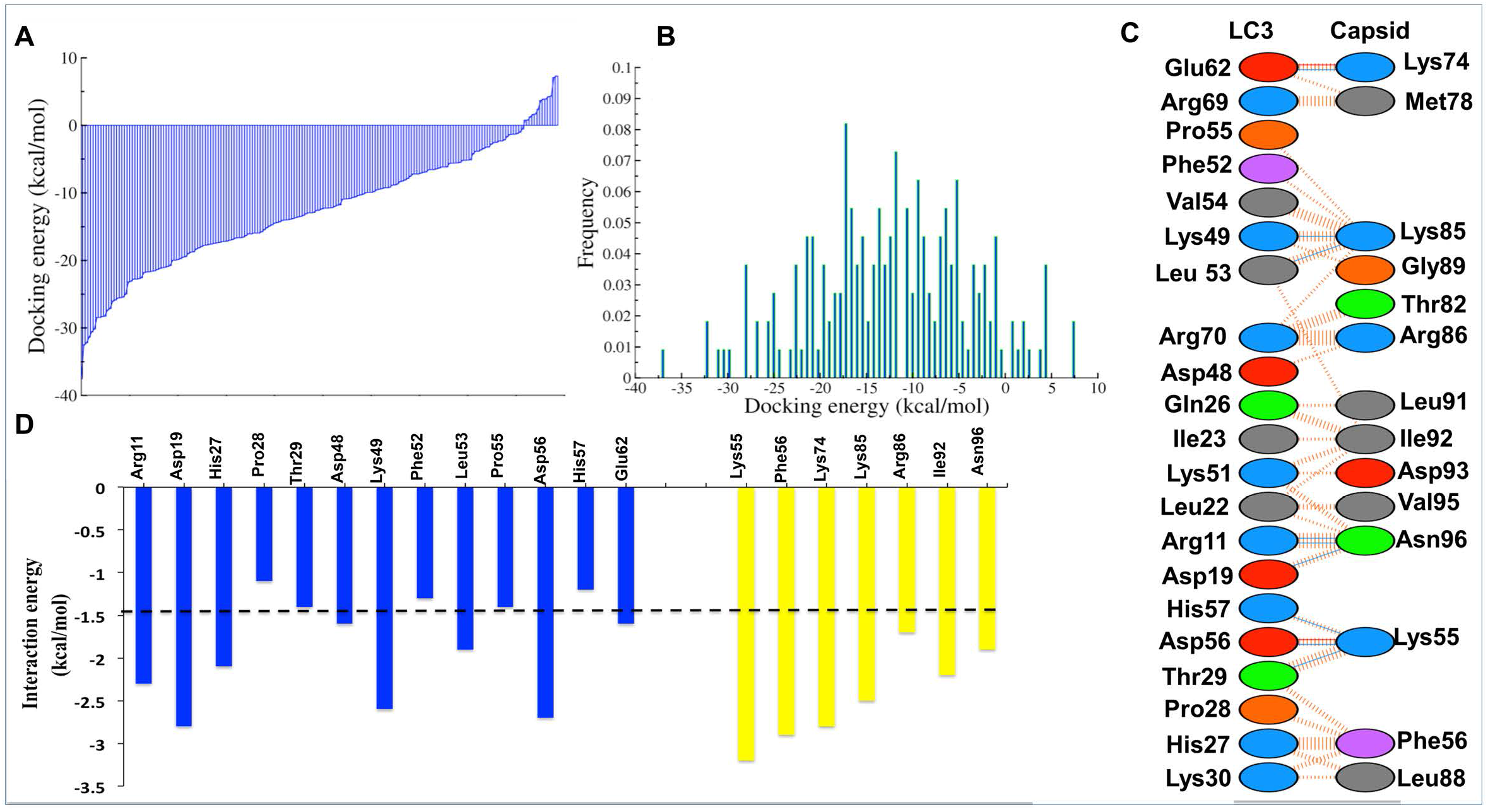
Quantitative analysis of capsid-LC3 interaction. (A) The docking energy values in kcal/mol, (B) Histogram of docking energy (kcal/mol), (C) Types of interactions between capsid (right) and LC3 (left). The hydrogen bonds are shown in direct blue lines and dotted orange lines shown the HpH contacts. The amino acid resides are marked as: basic (blue), acidic (red), aromatic (purple), polar (green) and HpH (grey). (D) The per-residue interaction was calculated for key residues. The cut-off (<−1.5 kcal/mol) is shown by dotted line.

## Discussion

Flavivirus replication complexes are 70-100 nm ER derived vesicles and convoluted membranes composed of viral NS proteins-NS1, NS2b, NS3, NS4a & NS5, each playing a crucial and independent role in the replication process. This membrane scaffold is also enriched in ER-resident proteins and utilizes a dynamic lipid-based sorting mechanism [10, 45–48].

Previous studies from our laboratory have mapped the immunofluorescence staining profile of several JEV NS proteins, E protein and ds RNA replication intermediate in infected MEFs, HeLa and Huh7 cells [11, 12, 27]. While the E protein is seen predominantly in the bulk ER-fraction and colocalizes with ER markers such as GRP78 [12], the NS proteins and dsRNA segregate in distinct punctate vesicles and membranes that show extensive overlap with the ERAD protein EDEM1 and LC3-I [27]. These represent sites of virus replication, and siRNA mediated depletion of LC3 (A&B) and of the ERAD proteins EDEM1 and Sel1L reduced JEV replication significantly highlighting the essential role of the ERAD proteins in the virus life-cycle [27]. Here we have characterized the localization pattern of capsid protein in JEV infected cells and observed its close association with LC3-I.

The capsid proteins of *flaviviruses*, including JEV have been reported to localize to both the cytoplasm and the nucleus (specifically the nucleolus) [49–51]. For JEV, the Gly42 and Pro43 were shown to be essential for nuclear localization of the JEV capsid protein [49]. These residues are highly conserved among the *flaviviruses*. The capsid protein has been shown to interact with LDs, and this interaction is crucial for virus replication and pathogenesis [17–22].

Similar to previous studies we also observed large foci of capsid protein in infected cells that were closely associated with replication complexes marked by NS1. While confocal microscopy images often showed extensive overlap between NS1 and capsid, these could be seen as close but distinct structures in high-resolution SIM images. The capsid also showed extensive localization on LDs, and quantitation of LD number showed a significant decrease in infected cells. However, studies with DENV and HCV have shown that LDs increase during infection [20, 52]. It is possible that lipid metabolism is differentially regulated by JEV which is primarily a neurotropic virus. Interestingly, a recent study with Poliovirus also shows a decrease in the LD number in infected cells [53].

It is now well appreciated that several positive strand RNA viruses utilize autophagy independent non-lipidated LC3 as a part of their replication scaffolds [27–31]. It is likely that these membrane supply topological platforms for segregation and protection of the virus replication machinery. The MHV, EAV and JEV replication complexes acquire the LC3 from the ERAD pathway and also contain additional ERAD proteins EDEM1 and SEL1L [27, 29, 30].

Here we have observed a strong association between JEV-capsid and LC3-I, both by high-resolution imaging and immunoprecipitation experiments. This association was also seen on nuclear localized capsid and LDs, and was independent of the autophagy receptor p62/SQSTM1. Capsid also did not show any association with the GABARAP family suggesting that this interaction is specific to LC3. These evidences suggest that there could be a direct interaction between capsid and LC3.

To gain further insights we have performed high resolution protein-protein docking studies between capsid and LC3. A putative LIR motif in the capsid protein was identified: **_56_FTAL_59_**, with the Phe56 and Leu59 residues showing high conservation across all flaviviruses. The C-terminal region of capsid that corresponds to the 4^th^ alpha-helix was also mapped to the LC3 interacting zone. Interestingly, several of the capsid residues that were identified in our modelling study: Met78, Lys 85, Arg86, Leu88, Leu91, Val95, Asn96 are significantly conserved between similar properties of amino acids in *flaviviruses*, with Lys85 and Arg86 being crucial for viral RNA packaging [19, 43].

LC3 though being a key component of the autophagy process also has several autophagy independent functions including those in virus replication [29, 30, 32, 54]. Studies have suggested that LC3 could serve as a source of membranes for efficient virus replication [28, 31]. We have also shown that siRNA mediated depletion of LC3 (A&B) significantly inhibits virus replication and egress [27]. The relevance of the capsid-LC3 association in the context of the JEV life-cycle needs further exploration. It is tempting to speculate that LC3 maybe part of an RNA exit channel for transport of the viral RNA from the replication compartment to a spatially distinct environment for nucleocapsid packaging. Indeed recent studies have shown an enrichment of RNA binding proteins in the LC3 dependent secretome interactome [55]. Further experiments to test the significance of the capsid-LC3 interaction in the nucleus and on LDs, and functional validation of our modelling data are subjects for our future studies.

### Materials and Methods Cell lines and virus

Huh7, HEK293T & C6/36 cells were obtained from the Cell Repository at the National Centre for Cell Sciences, Pune, India. HeLa cell line (CCL-2) was obtained from ATCC. WT and *Atg5-*deficient (*atg5*−/−) MEFs were a kind gift from Prof Noboru Mizushima and obtained through RIKEN Bio-Resource Cell Bank (RCB2710 and RCB2711). Huh7, HEK293T and MEFs were grown in Dulbecco’s modified Eagle’s medium (DMEM) (Himedia) supplemented with 10 % fetal bovine serum (FBS), 100 μg/ml penicillin/streptomycin, 2 mM L-glutamine and 1X MEM Non-Essential Amino Acids Solution (ACL006, Himedia). For all infection experiments, JEV isolate P20778 grown in C6/36 cells was used.

### Antibodies, reagents and plasmids

The following antibodies were used in the study: LC3 (Abcam: ab51520; Santa Cruz Biotechnology: sc-16756), JEV-NS1 (ab41651), JEV-Capsid (Genetex: GTX634152, GTX131368), p62/SQSTM1 (ab56416), GABARAP (ab109364), Myc-tag (ab9106; sc-70469), HA-tag (sc-53516; Sigma: H-6908), Mouse IgG (sc-2025), Rabbit IgG (Cell Signaling Technology: 2729S). Fluorescently labeled anti-mouse (A-11004, A-21202, A-21235), anti-rabbit (A-11008, A-11011, A-31573), and anti-goat antibodies (A-21469) secondary antibodies, BODIPY 493/503 (D3922) and ProLong Gold anti-fade reagent with DAPI (P36935) were obtained from Invitrogen, Thermo Fisher SCIENTIFIC. HRP-conjugated secondary antibodies were obtained from Jackson ImmunoResearch Laboratories Inc. The plasmids pCI-neo-myc-LC3 (deltaC22) (#45448) and pCI-neo-myc-LC3 (deltaC22, G120A) (#45449) were obtained from Addgene (deposited by Tamotsu Yoshimori) [33].

### JEV infection

All cells were mock- or JEV infected at 5 MOI (Huh7, MEFs) or 20 MOI (transfected HEK293T) for 24 h. Cells were then processed for immunostaining or immunoprecipiation and western blotting experiments.

### Plasmid Transfections

HEK293T cells were transfected with Myc-LC3δC22 or Myc-LC3δC22, G120A and 24 hpt, the cells were washed with PBS and infected with JEV. Cells were processed 24 hpt for immunostaining or immunoprecipiation and western blotting experiments. Transfections were done using TransIT^®^-LT1 Transfection Reagent (MIR2300) according to manufacturer’s protocol.

### Immunoprecipitation and Western blots

Mock/JEV-infected cells were lysed using 0.5 ml lysis buffer (20 mM Tris HCl, 1 mM EDTA, 250 mM, NaCl, 1% Triton X-100, 1 mM PMSF, 1 mM protease inhibitor). The total cell lysate was pre-cleared with protein A/G UltraLink^®^ resin (53132). 5ug of the specific immunoprecipitating antibody was added to the pre-cleared lysate: LC3 (ab51520); Capsid (GTX131368); myc-tag (sc-70469) along with specific IgG controls: rabbit IgG (2729S) and mouse IgG (sc-2025). Lysate-antibody complex was allowed to interact for 8 h at 4°C, followed by immobilization with protein A/G UltraLink^®^ resin. Bound proteins from the beads were eluted using 2X SDS sample buffer and analyzed by gel electrophoresis and western blotting. Each immunoprecipitation experiment was performed two or more times.

### Immunostaining, fluorescence microscopy & image processing

For immunofluorescence experiments, cells were seeded on poly-L-lysine coated coverslips. For overexpression and colocalization studies, the cells were transfected (HEK293T) and mock- or JEV infected as described. Each experiment had biological duplicates, and was performed three or more times. Following transfection or infection, cells were fixed with 2% paraformaldehyde and permeabilized with 0.3% Tween-20 for 30 min at RT. Blocking is done with 1% Bovine serum albumin (BSA; Sigma, A7906) in PBS for 1 h at RT prior to incubation with primary antibody. The cells were washed thrice with 1% BSA for 15 min and then stained with Alexa Fluor labeled specific secondary antibodies for 1 h at RT. After washing, the coverslips were mounted on ProLong Gold anti-fade reagent with DAPI. For lipid droplet staining, cells were incubated with BODIPY 493/503 (1ug/ml) along with the primary antibodies. All antibodies and BODIPY 493/503 were diluted in the blocking solution. Images were acquired on an Olympus FV3000 confocal microscope with 60X (NA 1.4) objective. For colocalization experiments, Z-stacks were acquired at 0.41 μm per slice by sequential scanning with a 60X objective lens. The colocalization analysis and LD quantification (ALDQ) was done using ImageJ software [56]. SIM images were acquired on the Deltavision OMX SR imaging system, Cytiva (formerly GE). All immunofluorescence experiments were performed in biological duplicates. Images shown are representative of two or more independent experiments. Pearson coefficient calculations were from two or more independent experiments.

### Computational modelling & docking studies

#### System Preparation and validation

A detailed molecular modelling was performed for the JEV-capsid (PDB-ID: 5OW2, UniProt ID: E7CG11) and LC3 (PDB-ID: 1UGM, UniProt ID: Q62625) proteins. The capsid dimer was generated through homology modelling. The structures were optimized and then minimized using the Protein Preparation Wizard module of Maestro (Schrödinger Release 2020-1: Maestro, Schrödinger, LLC, New York, NY, 2020 [57, 58]. Since the complete active form models of capsid and LC3 are not available, homology models were generated through the modeller [59] to understand the complete structures. The robustness of predicted model structure was assessed using various validation servers such as PROCHECK [60, 61] and ProSA-Web (Z-score) [61].

#### Protein -protein docking study

Molecular dynamics minimization was employed to allow conformational relaxation of the protein structures prior to subjecting them to protein–protein docking calculations [58]. Subsequently, two different algorithms (PyDOCK & Swarmdock) were used to perform the protein-protein dockings to identify the most likely binding interfaces and poses of capsid-LC3 interactions. The docking procedure aimed to generate a set of solutions for candidates with at least one near native structure. Since, rigid docking through PyDock [62] allows some steric clashes, flexible docking by Swarmdock [63, 64] was also done on relaxed structures of the proteins. The candidate solutions were scored and ranked according to different parameters such as clusters, lowest binding energy, number of conformers, and agreement with known binding sites. Clustering of docked poses was conducted to filter out the most likely complex of capsid-LC3. We quantified the docking results and picked the lowest energy zone (< −30.0 kcal/mol, Fig 8A-B). Only the best-docked pose, which ranged between <−30.0 to −40.0 kcal/mol, was used for further analysis (Fig 8C-D).

## Acknowledgements

We gratefully acknowledge Nisheeth Agarwal, Sankar Bhattacharyya and Amitabha Mukhopadhyay for inputs and suggestions, and all members of the Molecular Virology labs at THSTI & RCB.

